# Intestinal inflammation and altered gut microbiota associated with inflammatory bowel disease renders mice susceptible to *Clostridioides difficile* colonization and infection

**DOI:** 10.1101/2020.07.30.230094

**Authors:** Lisa Abernathy-Close, Madeline R. Barron, James M. George, Michael G. Dieterle, Kimberly C. Vendrov, Ingrid I. Bergin, Vincent B. Young

## Abstract

*Clostridioides difficile* has emerged as a noteworthy pathogen in patients with inflammatory bowel disease (IBD). Concurrent IBD and CDI is associated with increased morbidity and mortality compared to CDI alone. IBD is associated with alterations of the gut microbiota, an important mediator of colonization resistance to *C. difficile*. Here, we describe and utilize a mouse model to explore the role of intestinal inflammation in susceptibility to *C. difficile* colonization and subsequent disease severity in animals with underlying IBD. *Helicobacter hepaticus*, a normal member of the mouse gut microbiota, was used to trigger inflammation in the distal intestine akin to human IBD in mice that lack intact IL-10 signaling. Development of IBD resulted in a distinct intestinal microbiota community compared to non-IBD controls. We demonstrate that in this murine model, IBD was sufficient to render mice susceptible to *C. difficile* colonization. Mice with IBD were persistently colonized by *C. difficile*, while genetically identical non-IBD controls were resistant to *C. difficile* colonization. Concomitant IBD and CDI was associated with significantly worse disease than unaccompanied IBD. IL-10-deficient mice maintained gut microbial diversity and colonization resistance to *C. difficile* in experiments utilizing an isogenic mutant of *H. hepaticus* that does not trigger intestinal inflammation. These studies in mice demonstrate that the IBD-induced microbiota is sufficient for *C. difficile* colonization and that this mouse model requires intestinal inflammation for inducing susceptibility to CDI in the absence of other perturbations, such as antibiotic treatment.

**IMPORTANCE:** The incidence of CDI continues to increase significantly among patients with IBD, independent of antibiotic use, yet the relationship between IBD and increased risk for CDI remains to be understood. However, antibiotic-induced perturbations of the gut microbiota may mask mechanisms specific to IBD-induced *C. difficile* susceptibility and infection. Our study sought to describe and utilize a mouse model to specifically explore the relationship between the IBD-induced gut microbial community and susceptibility to *C. difficile* colonization and CDI development. We demonstrate that IBD is sufficient for *C. difficile* colonization and infection in mice and results in significantly worse disease than IBD alone, representing a murine model that recapitulates human IBD and CDI comorbidity. Furthermore, this model requires IBD-induced inflammation to sculpt a microbiota permissible to *C. difficile* colonization. Use of this model will aid in developing new clinical approaches to predict, diagnose, and treat *C. difficile* infection in the IBD population.

## INTRODUCTION

Inflammatory bowel diseases, including Crohn’s disease and ulcerative colitis, are chronic and progressive diseases characterized by inflammation of the digestive tract. The incidence of *C. difficile* infection (CDI) has significantly increased among hospitalized patients with inflammatory bowel disease (IBD) over the past two decades (1–3). *C. difficile* is a spore-forming bacterium that produces enterotoxins that damage the intestinal epithelium. *C. difficile* was initially described as the as a cause of antibiotic-associated diarrhea (4), highlighting the role of the gut microbiota in CDI disease pathogenesis. Normally, an intact intestinal microbiota provides colonization resistance to *C. difficile* colonization and subsequent infection (5). However, antibiotic exposure can render otherwise healthy individuals susceptible to *C. difficile* colonization and CDI disease due to disruption of the microbiota. Although antibiotic use is a known risk factor for CDI, other risk factors have been recognized, including immunosuppression and pre-existing IBD (6, 7). Alterations in the gut microbiota are known to occur in patients with IBD (8, 9), independent of antimicrobial exposure. CDI is associated with more severe intestinal microbiota disturbances among patients with IBD (10). In addition, the efficacy of fecal microbiota transplantation (FMT) to treat CDI and the presence of particular microbial taxa is affected by underlying IBD (11), indicating that the pathophysiology of IBD influences gut microbiota composition and CDI outcomes.

Various aspects of IBD and CDI have long been studied in separate animal models (12, 13), yet a robust mouse model of comorbid IBD and CDI in the absence of antibiotic-induced *C. difficile* colonization perturbation of the microbiota has yet to be described. *Helicobacter hepaticus* colonization in mice genetically predisposed to developing colitis, such as those lacking the regulatory cytokine IL-10, are useful research tools that mimic human inflammatory bowel disease processes (14–16). IL-10-deficient mice reared in specific pathogen-free (SPF) conditions develop colitis that resembles human IBD when colonized with *H. hepaticus* (15, 17, 18) and this intestinal inflammation is associated with alterations in gut microbiota community structure (19). Interestingly, the ability for *H. hepaticus* to trigger IBD in IL-10-deficient mice depends on the expression of cytolethal distending toxin and the presence of an indigenous microbiota, as germ-free mice or mice colonized with CDT-deficient *H. hepaticus* do not develop intestinal inflammation (20, 21). Most previously described mouse models of CDI require antibiotic administration to disrupt the intestinal microbiota and render animals susceptible to *C. difficile* colonization and disease (13, 22), and have revealed that colonization resistance and protection from CDI is mediated by the microbiota and host immune responses (5, 23).

In the present study, we sought to study the specific relationship between IBD and CDI concurrently in a mouse model. We wished to develop a system where we could evaluate the role of intestinal inflammation in the induction of susceptibility to *C. difficile* colonization. These results demonstrate that mice with IBD harbor an altered microbiota, compared to mice without IBD, that renders animals susceptible to *C. difficile* colonization and infection in the absence of antibiotic treatment.

## MATERIALS AND METHODS

### Mice

Male and female C57BL/6 wild-type or IL-10-deficient mice were maintained under specific pathogen-free (SPF), *Helicobacter*-free conditions. Mice were at least 8 weeks of age mice at the start of experiments. All mice were from a breeding colony at the University of Michigan that were originally derived from Jackson Laboratories almost 20 years ago. Euthanasia was carried out via CO_2_ inhalation at the conclusion of the experiment. Animal studies were approved by The University of Michigan Committee on the Care and Use of Animals animal husbandry was performed in an AAALAC-accredited facility.

### *Helicobacter hepaticus* strains, growth conditions, and murine inoculation

*H. hepaticus* strain 3B1 (ATCC 51488) was obtained from the American Type Culture Collection (Manassas, VA). The isogenic mutant 3B1::Tn20 has a transposon inserted near the start of *cdtA* and no longer produces cytolethal distending toxin (CDT) (20). Wild-type *H. hepaticus* 3B1 and 3B1::Tn20 were grown on tryptic soy agar (TSA) supplemented with 5% sheep blood at 37°C for 3 to 4 days in a microaerobic chamber (1-2% oxygen, Coy Laboratories). The isogenic mutant 3B1::Tn20 is chloramphenicol-resistant and was grown on media additionally supplemented with 20 μg/ml chloramphenicol (Sigma, St. Louis, MO). *H. hepaticus* suspensions for animal inoculation were prepared by harvesting organisms from culture plates into trypticase soy broth (TSB). Mice were challenged with 10^8^ CFU *H. hepaticus* by oral gavage. *H. hepaticus* colonization status in mice and colonizing strain were confirmed by PCR of the *cdtA* gene (24) on fecal DNA prior to *C. difficile* spore challenge.

### *C. difficile* strain and growth conditions

The *C. difficile* reference strain VPI 10463 (ATCC 43255) was used as previously described in a murine model of CDI by Theriot *et al.* (22). To determine a correct dose of *C. difficile* spores per challenge, viable spores in each inoculum were enumerated by plating for colony-forming units (CFU) per mL on pre-reduced taurocholate cycloserine cefoxitin fructose agar (TCCFA). TCCFA was prepared as previously described (25). TCCFA pates with fecal or cecal samples or spore inoculum were incubated in an anaerobic chamber (Coy Industries) at 37°C for 18 hours prior to colony enumeration.

### Antibiotic administration

Mice were rendered susceptible to *C. difficile* infection by treating mice with 0.5 mg/mL cefoperazone (MP Pharmaceuticals) in sterile distilled drinking water (Gibco) ad libitum. The antibiotic-supplemented water was provided for 10 days, followed by 2 days of drinking water without antibiotics (22).

### *C. difficile* spore challenge

After challenge with *H. hepaticus*, TSB vehicle, or antibiotic pretreatment, animals were then challenged by oral gavage with 10^3^ – 10^4^ CFU *C. difficile* spores suspended in 50 µl of distilled water (Gibco) or mock-challenged with water vehicle. Over the course of the experiment, mice were regularly weighed and feces were collected for quantitative culture.

### *C. difficile* quantification

Fresh feces were collected from each mouse into a pre-weighed sterile tube. Immediately following collection, the tubes were re-weighed to determine fecal weight and passed into an anaerobic chamber (Coy Laboratories). Each sample was then diluted 10% (w/v) with pre-reduced sterile phosphate buffered saline (PBS) and serially diluted onto pre-reduced TCCFA plates with or without erythromycin supplementation. The plates were incubated anaerobically at 37°C, and colonies were enumerated after 18 to 24 hours of incubation.

### Quantitative detection of *C. difficile* toxin in cecal contents

Functional *C. difficile* toxin was measured using a real-time cellular analysis (RTCA) assay (26). The RTCA assay was used to detect changes in cell-induced electrical impedance in cultured colorectal cell monolayers in response to cecal contents collected from mice with CDI to determine concentrations of active toxin. Cecal contents collected from mice at the time of euthanasia were weighed and brought to a final dilution of 1:1000 (w/v) with sterile 1X PBS. Diluted cecal contents were allowed to settle in the original collection tubes prior to transferring supernatant aliquots to fresh tubes. Cecal content supernatants were then filtered through a sterile 0.22 μm 96-well filter plates (E-Plate VIEW 96, ACEA Biosciences) and plates were centrifuged at 5000 x g for 10 minutes at room temperature. HT-29 cells, a human colorectal adenocarcinoma cell line with epithelial morphology (ATCC HTB-38), were seeded in 96 well plates in Dulbecco’s Modified Eagle Media (DMEM) and allowed to grow to a confluent monolayer overnight prior to loading processed cecal content supernatant or purified *C. difficile* toxin A (List Biological Labs). Samples were run in triplicate. A standard curve was generated using wells that received purified *C. difficile* toxin A; this also served as a positive control. Prior to adding the sample to the plates containing HT29 monolayers, an aliquot of each sample, also run in triplicate, was incubated in parallel with anti-toxin specific for *C. difficile* toxin A and B (TechLab, Blacksburg, VA) for 40 minutes at room temperature to confirm the presence of *C. difficile* toxin in samples by neutralizing the cytotoxic activity. Active *C. difficile* toxin causes cytotoxic effects on the HT29 cells, which results in a dose-dependent and time-dependent decrease in cell impedance. Cell impedance (CI) data following incubation with cecal contents from mice with CDI was acquired and analyzed using the xCELLigence RTCA system and software (ACEA Biosciences, San Diego, CA). A normalized CI was calculated for each sample by normalizing the CI to the last CI measured at the time point prior to adding cecal content to the well.

### Clinical disease severity and histopathological damage scoring

Mice were monitored for clinical signs of disease. Disease scores were averaged based on scoring of the following features for signs of disease: weight loss, activity, posture, coat, diarrhea, eyes/nose. A 4-point scale was assigned to score each feature and the sum of these scores determined the clinical disease severity score (27). Formalin-fixed tissue sections prepared from cecum and colon were H&E stained and evaluated by a blinded animal pathologist. Histopathologic damage in each tissue was scored using epithelial destruction, immune cell infiltration, and edema on a 4-point scale for each category and the sum of these scores determined the histological score (22, 25, 28).

### DNA extraction and 16S rRNA gene sequencing

Cecal and colon luminal contents were separately collected from mice with IBD and without IBD at the time point immediately preceding *C. difficile* spore challenge. The University of Michigan Microbiome Core extracted total DNA from cecal and colon contents and prepped DNA libraries as previously described (29). The V4 region of the 16S rRNA gene was amplified from each sample using the dual indexing sequencing strategy as described previously (30). Sequencing was done on the Illumina MiSeq platform using the MiSeq Reagent kit V2 (#MS-102-2003) to sequence the amplicons (500 total cycles) with modifications found in the Schloss SOP (https://github.com/SchlossLab/MiSeq_WetLab_SOP). The V4 region of the mock community (ZymoBIOMICS Microbial Community DNA Standard, Zymo Research) was also sequenced to supervise sequencing error. Data were analyzed using mothur (v 1.42.3) (31).

### Statistics

Statistical analysis using unpaired student’s t-test or one-way analysis of variance (ANOVA) with Tukey’s post-hoc test was performed using R. Clinical scores were analyzed using the Mann-Whitney U test. A p-value less than 0.05 was considered statistically significant.

### Data availability

Code and processing information are available on GitHub repository: https://github.com/AbernathyClose/AbernathyClose_IbdCdi_mBio_2020.

## RESULTS

### Intestinal inflammation in IL-10-deficient mice colonized with H. hepaticus is associated with altered gut microbiota

Wild-type (WT) and IL-10-deficient C57BL/6 mice reared under specific pathogen-free (SPF) conditions were colonized with *H. hepaticus* or gavaged with vehicle and subsequently evaluated for intestinal inflammation and gut microbiota diversity. We sought to confirm that wild-type mice reared under SPF conditions do not develop signs of intestinal inflammation regardless of *H. hepaticus* colonization status (Figure 1A and 1B). We found that intestinal inflammation in IL-10-deficient mice was observed at 7-14 days after IBD was triggered by *H. hepaticus* infection. The level of the inflammatory marker lipocalin-2 was significantly increased in feces 7 days after *H. hepaticus* colonization, and this increase was sustained at 14 days post-colonization (Figure 1A). WT mice did not have increased levels of fecal lipocalin-2, regardless of *H. hepaticus* colonization status (Figure 1A). Histological examination of colon sections harvested from and IL-10-deficient mice revealed pathology consistent with inflammatory bowel disease, including loss of goblet cells, inflammatory cell infiltration, and crypt elongation at 14 days post-*H. hepaticus* colonization, compared to WT mice or either genotype receiving vehicle (Figure 1B).

**Figure 1.**
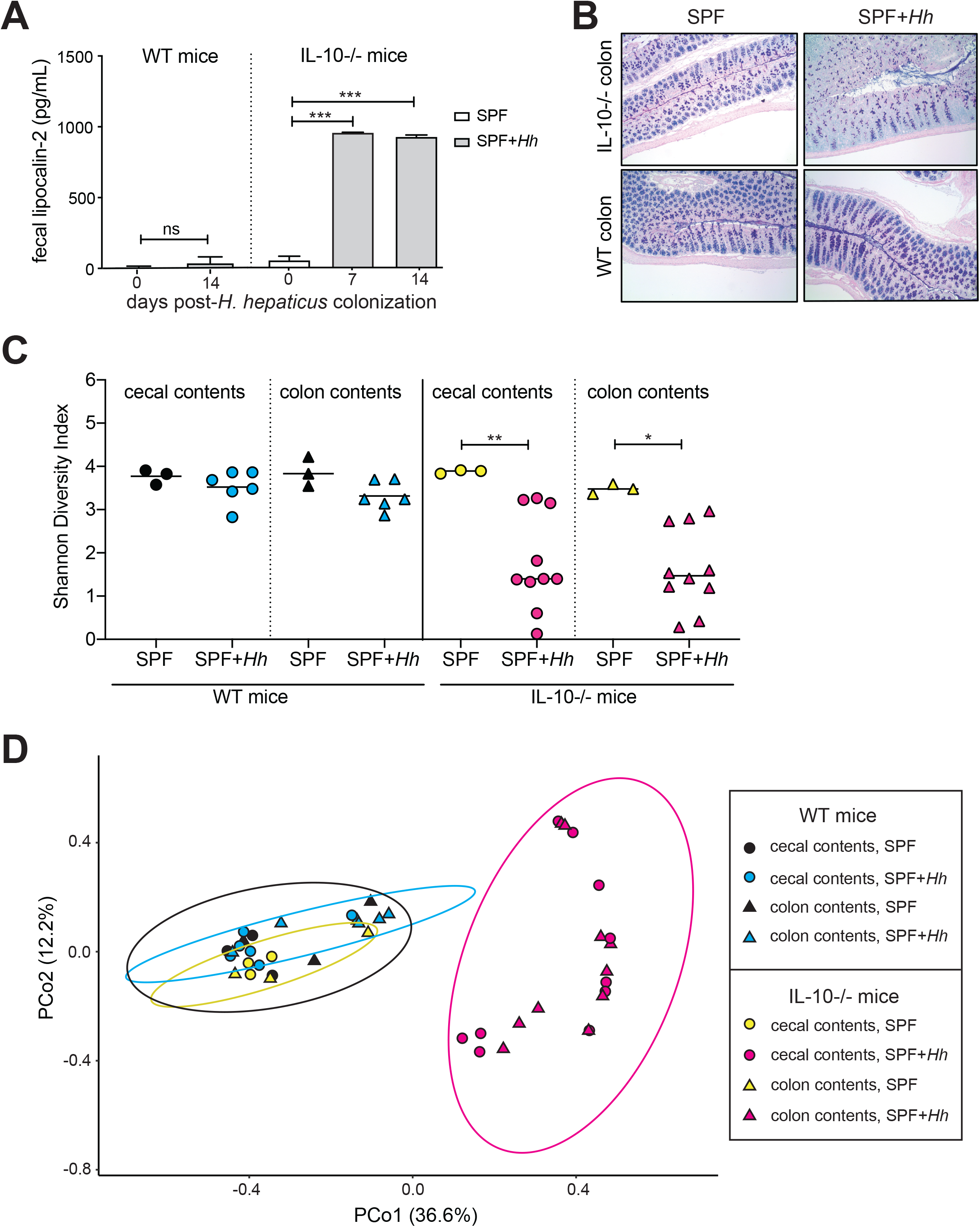
Intestinal inflammation is associated with altered intestinal microbiota in mice. A) Lipocalin-2 levels in feces from SPF wild-type (WT) mice or SPF IL-10−/− mice were measured by ELISA at day 7 and 14 post-colonization with *H. hepaticus* (*Hh*) or mock-challenged with vehicle. ANOVA and Tukey test, ***p<0.001. n = 6-13 mice/group. B) Colonic mucin and mucosal integrity in SPF WT mice or SPF IL-10−/− mice 14 post-colonization with *H. hepaticus* or mock-challenged with vehicle were visualized with alcian blue/periodic acid-Schiff staining of colon tissue sections (representative images, x100 magnification). C) Shannon diversity index of luminal contents collected from the cecum (round) and colon (triangle) of SPF WT mice or SPF IL-10−/− mice at 14 days post-*H. hepaticus (Hh)* colonization or mock-challenged with vehicle. Two-tailed unpaired t-test, *p < 0.05. D) Principal coordinates analysis plot of Bray-Curtis distances of luminal content collected from the cecum (round) and colon (triangle) of SPF WT mice or SPF IL-10−/− mice 14 days post-*H. hepaticus (Hh)* colonization or mock-challenged with vehicle. 95% confidence ellipses are shown. SPF = specific pathogen-free, *Hh* = *Helicobacter hepaticus*.

We then sought to determine if intestinal inflammation was associated with changes in gut microbiota diversity. *H. hepaticus* colonization in WT mice does not induce IBD and thus does not significantly impact diversity of the distal gut microbial community (Figure 1C and 1D). However, intestinal inflammation induced by *H. hepaticus* colonization in SPF IL-10-deficient was associated with altered microbial community structure (Supplemental Figure 1) and significantly less diversity microbiota (Figure 1C and 1D), compared to non-inflamed counterparts. Taken together, these data demonstrate that mice with IBD harbor altered gut microbiota compared to mice without IBD, and that the presence of intestinal inflammation is associated with significant alterations in alpha and beta diversity of the distal gut microbiota.

### Altered intestinal microbiota associated with IBD is sufficient to induce susceptibility to C. difficile colonization

We explored whether mice with intestinal inflammation due to active IBD are susceptible to *C. difficile* colonization and CDI disease, and compared these outcomes to those following antibiotic pretreatment in mice lacking IBD. SPF IL-10-deficient mice were either treated with the broad-spectrum antibiotic cefoperazone or colonized with *H. hepaticus* to trigger IBD prior to *C. difficile* strain VPI 10463 spore challenge (Figure 2A) and monitored for *C. difficile* colonization and signs of clinical disease. Mice with established IBD were challenged with *C. difficile* spores and then monitored for *C. difficile* colonization. *C. difficile* colonization was determined by plating feces collected from mice for up to 7 days post-spore challenge on selective media and cecal contents were plated when mice were euthanized.

**Figure 2.**
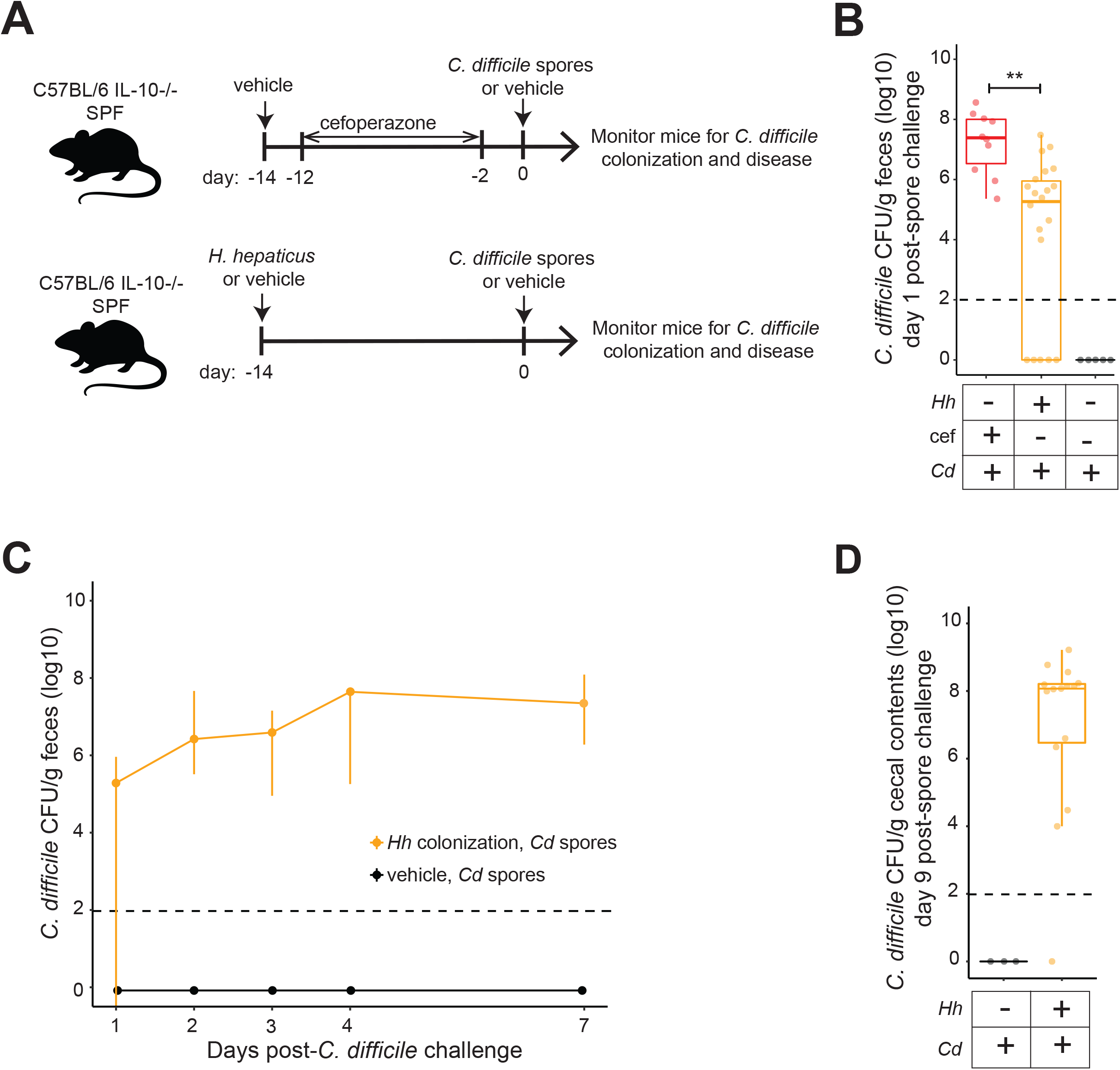
*C. difficile* colonization dynamics in mice with IBD differ compared to antibiotic-pretreated mice without IBD. A) Mouse model of CDI after antibiotic treatment or IBD triggered by *H. hepaticus* colonization. B) *C. difficile* burden in feces one day after *C. difficile* spore challenge. Feces were collected and plated anaerobically on selective agar plates to quantify *C. difficile* burden. Two-tailed unpaired t-test, **p < 0.01. C) Quantification of *C. difficile* in feces collected from mice with or without IBD for up to 7 days post-*C. difficile* spore challenge. Data are presented as median and interquartile range for each time point. D) *C. difficile* colonization level in cecal contents harvest from mice with and without IBD at 9 days post-*C. difficile* spore challenge. Data represent 2-3 independent experiments and are plotted as mean +/− standard deviation unless otherwise indicated. Dotted line indicates limit of detection for *C. difficile* quantification (10^2^ CFU).

IL-10-deficient mice challenged with *C. difficile* spores in the absence of microbiota perturbation due to IBD or antibiotic pretreatment were resistant to *C. difficile* colonization (Figure 2B). Conversely, cefoperazone-treated mice had high levels of *C. difficile* colonization one day post-spore challenge, while mice with IBD had significantly lower levels of *C. difficile* colonization at this time point (Figure 2B). We found that while 100% of mice treated with cefoperazone and subsequently challenged with *C. difficile* spores were colonized by day one post-spore challenge, only 68% of mice with IBD had detectable levels of *C. difficile* in feces at this time point. However, 89% of mice with IBD were colonized 7 days post-spore challenge, and one mouse remained colonization resistant at 9 days after spore challenge, lacking cultivatable *C. difficile* from cecal contents (Figure 2D). This experimental endpoint of day 9 post-*C. difficile* spore challenge was arbitrarily pre-determined in the event that mice with IBD did not develop severe disease resulting in a moribund condition upon *C. difficile* colonization. These experiments establish a mouse model of IBD rendering mice susceptible to *C. difficile* colonization, obviating the need for antibiotics for CDI development.

To follow up on the finding that mice with IBD are susceptible to *C. difficile* colonization, we assessed disease severity associated with concurrent IBD and CDI compared to antibiotic-induced CDI by monitoring weight loss, immune response, and scoring clinical disease. Mice with CDI following cefoperazone treatment lost a significant amount of weight a hallmark of severe disease associated with CDI, compared to mock or antibiotic-treated controls. Mice with concomitant IBD and CDI lost significantly more weight compared to mice with IBD alone at day 7 and day 9 post*-C. difficile* spore challenge (Figure 3A), indicating more severe disease associated with comorbid disease compared to unaccompanied IBD. Furthermore, mice with CDI associated with underlying IBD demonstrated significantly higher clinical disease scores compared to mice with IBD alone (Figure 3B). Neutrophil and eosinophil responses are associated with severe CDI (32–34). CDI occurring after cefoperazone treatment resulted in a significant increase in circulating neutrophils and eosinophils at day 2-3 post-*C. difficile* spore challenge, compared to antibiotics or spores alone (Figure 3C). CDI occurring after cefoperazone treatment resulted in a significant increase in circulating neutrophils and eosinophils at day 2 post-*C. difficile* spore challenge, compared to antibiotics or spores alone (Figure 3C). Comorbid IBD and CDI was associated with significantly less blood eosinophils compared to CDI following antibiotic treatment (Figure 3C). These data reveal that mice with underlying IBD persistently colonized with a toxigenic strain of *C. difficile* develop an altered course of disease associated with CDI, compared to their antibiotic pretreated counterparts.

**Figure 3.**
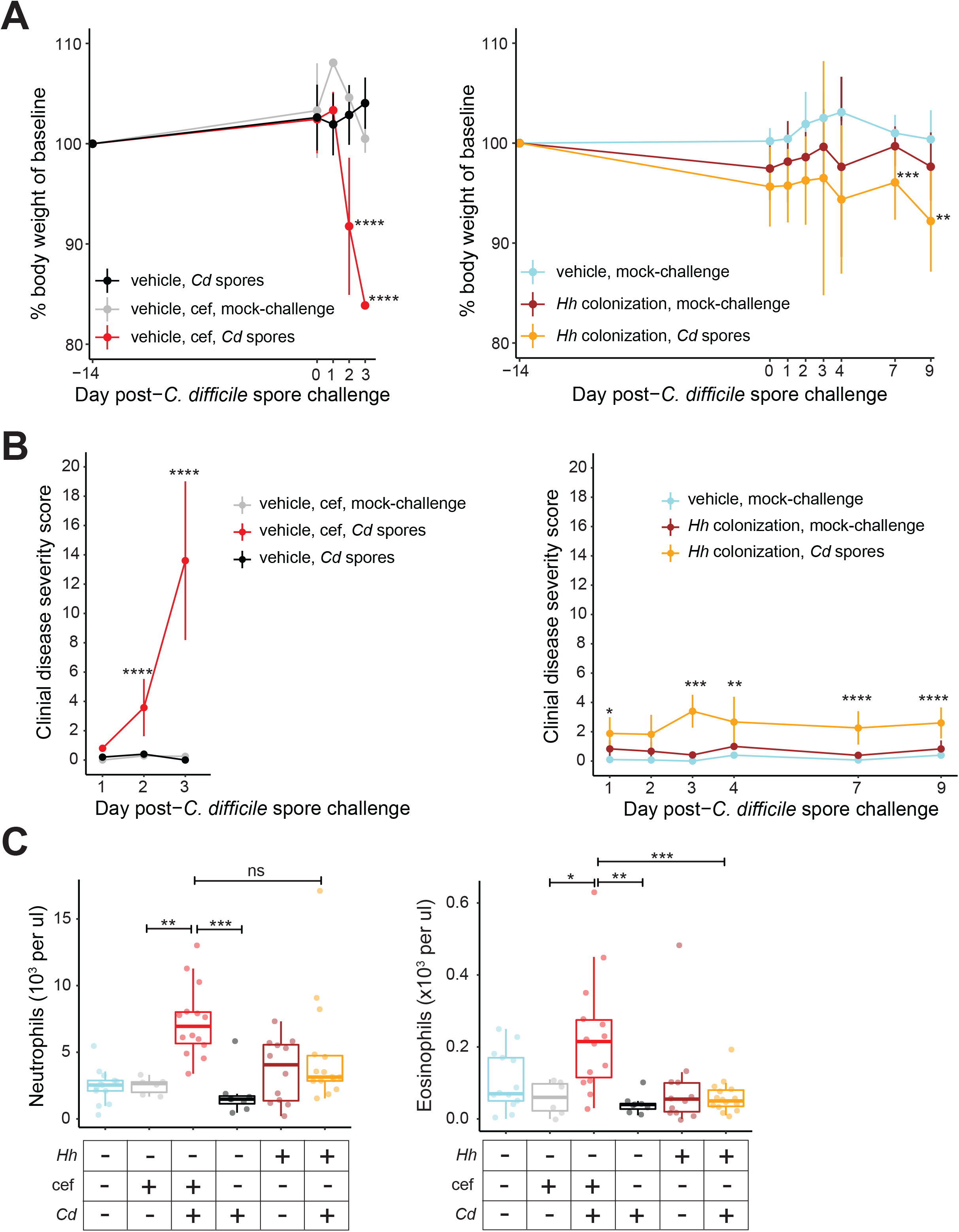
Mice with IBD do not develop severe CDI despite stable *C. difficile* colonization. A) Weight loss and B) clinical disease severity were assessed in mice with antibiotic-associated CDI or IBD, as well as concurrent IBD and CDI. C) Neutrophil and eosinophil levels in the blood at day 2 or day 9 post-CDI. Data represent 2-3 independent experiments and are plotted as mean +/− standard deviation and analyzed by ANOVA or Mann-Whitney test, *p > 0.05, **p > 0.01, ***p > 0.001, ****p > 0.0001. ns = not statistically significant.

### Toxigenic C. difficile produces significantly less toxin in cecum of mice with IBD compared to antibiotic-pretreated mice

We sought to further elucidate the cause of differences in clinical disease severity observed in antibiotic-versus IBD-associated CDI by examining intestinal pathology. *C. difficile* toxin mediates disease due to CDI by causing damage to the intestine (35, 36). Therefore, we investigated intestinal histopathology and corresponding functional *C. difficile* toxin levels in antibiotic pretreated mice with CDI (day 2 post-CDI) compared to comorbid IBD and CDI (day 9 post-CDI) (Figure 4). Histopathology of the cecum and colon of mice was quantified by a veterinary pathologist that scored tissues for the amount of edema, infiltration of leukocytes, and epithelial damage. We established that mice pretreated with cefoperazone had significantly higher cecum and colon histopathology scores, reflective of more intestinal pathology, while histopathology scores were similar between mice with IBD alone and mice with comorbid CDI (Figure 4A). Interestingly, despite similar cecal burden of *C. difficile* colonization (Figure 4B), mice with CDI following cefoperazone treatment had significantly more active *C. difficile* toxin in cecal contents compared to mice with comorbid IBD and CDI (Figure 4C). Furthermore, several mice with IBD colonized with *C. difficile* did not have detectable *C. difficile* toxin activity, in contrast to antibiotic pretreated mice with CDI (Figure 4C).

**Figure 4.**
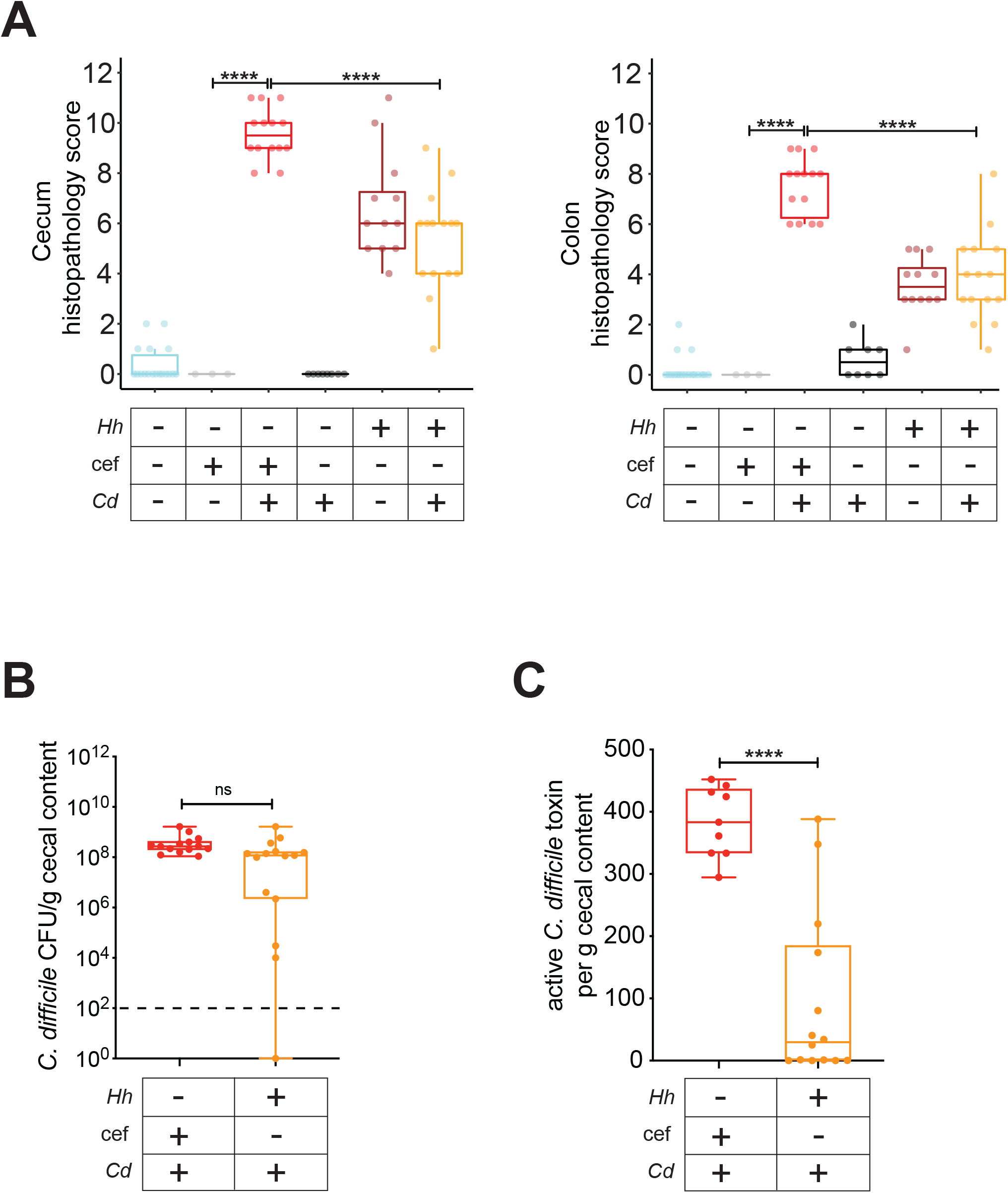
Mice pretreated with antibiotics have more intestinal damage and harbor more active *C. difficile* toxin in cecal contents compared to colonized mice with IBD. A) Histopathological damage in cecum and colon collected from mice with antibiotic- or IBD-associated CDI. Epithelial destruction, immune cell infiltration, and edema were scored on a 4-point scale for each category, and the sum of these scores determined the histological score in each tissue. B) *C. difficile* colonization burden in cecal contents at experimental endpoint (2 days post-spore challenge for cefoperazone pretreated mice and 9 days post-spore challenge for mice with IBD). C) Active *C. difficile* toxin level in cecal contents at experimental endpoint was quantified by real-time cell analysis (RTCA). Samples were also incubated with anti-toxin specific for *C. difficile* toxin A to confirm the presence of *C. difficile* toxin in toxin-positive samples by neutralizing the cytotoxic activity (data not shown). Data represent two independent experiments. Data represent two independent experiments and are plotted as mean +/− standard deviation and analyzed by ANOVA or Mann-Whitney test, ****p > 0.0001. ns = not statistically significant.

### Il-10-deficient mice colonized with a strain of H. hepaticus that lacks induction of intestinal inflammation are resistant to C. difficile colonization and infection

The mouse model of IBD and CDI we describe in the present study necessitates the use of *H. hepaticus* to trigger intestinal inflammation in mice lacking intact IL-10 signaling. Therefore, we utilized an isogenic strain of *H. hepaticus* that does not trigger IBD in IL-10-deficient mice (20, 21) to explore the contribution of *H. hepaticus* in the gut microbiota to subsequent *C. difficile* colonization and infection. IL-10-deficient mice reared in a SPF environment were colonized with either the wild-type strain *H. hepaticus* 3B1 (*Hh*CDT+) or a CDT-deficient strain *H. hepaticus* 3B1:Tn20 (*Hh*CDT−), and 14 days later mice were challenged with *C. difficile* strain 10463 spores and monitored for *C. difficile* colonization and disease (Figure 5A). We measured fecal lipocalin-2 levels by ELISA in SPF IL-10-deficient mice 14 days after colonization with *Hh*CDT+ or *Hh*CDT- and found that mice colonized with wild-type *H. hepaticus* had significantly more fecal lipocalin-2 than those colonized with CDT-deficient *H. hepaticus* (Figure 5B). Furthermore, intestinal inflammation triggered by wild-type *H. hepaticus* colonization was associated with lower microbiota diversity in luminal contents collected from cecum and colon, compared to non-inflamed mice colonized with CDT-deficient *H. hepaticus* (Figure 5C). These results confirm previous reports showing that CDT-deficient *H. hepaticus* does not trigger IBD in IL-10-deficient mice (20, 21), and demonstrate that this lack of intestinal inflammation is associated with higher distal gut microbial diversity compared to mice with wild-type H*. hepaticus*-induced IBD.

**Figure 5.**
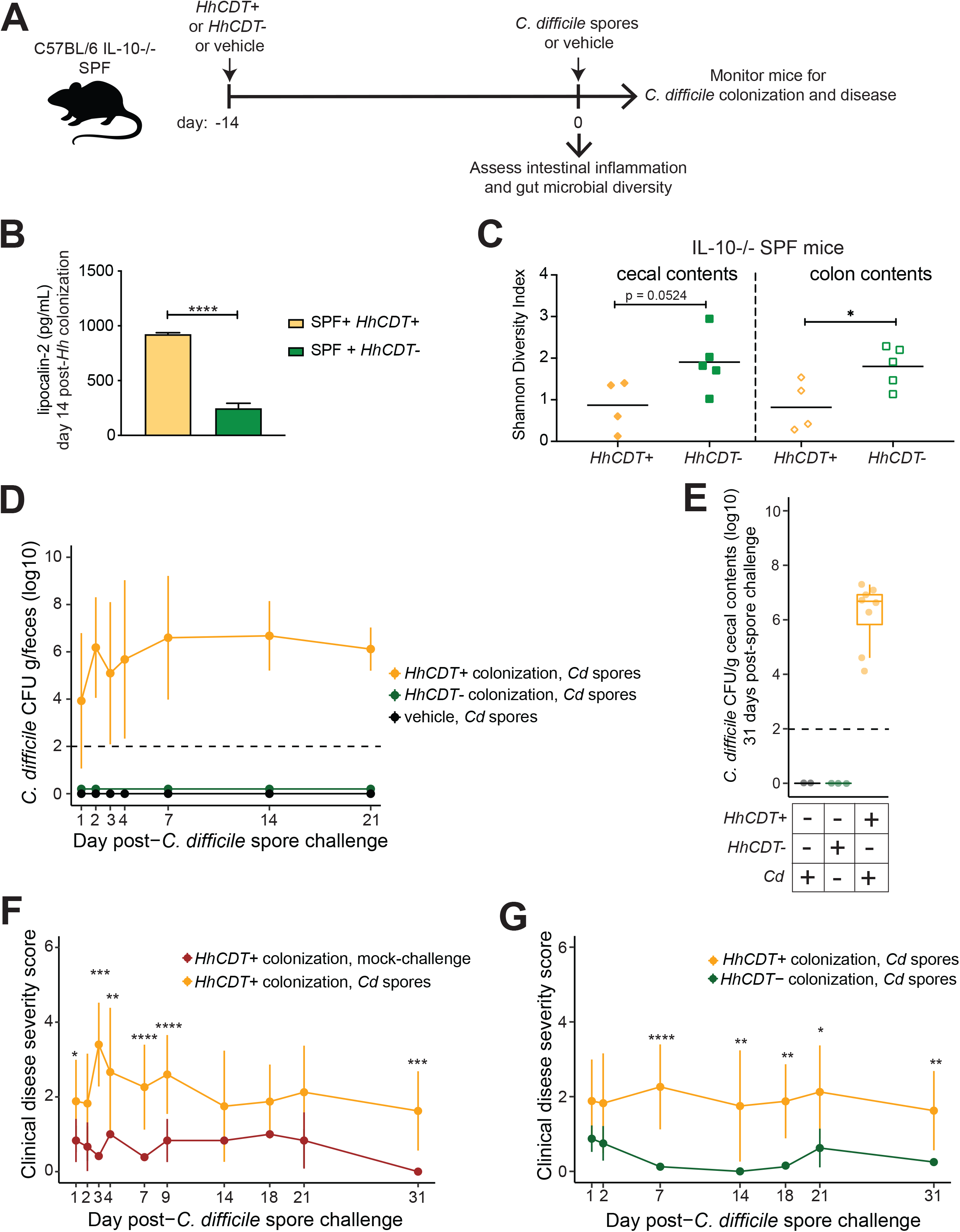
IL-10−/− mice colonized with isogenic strain of *H. hepaticus* that does not cause IBD are resistant to *C. difficile* colonization. A) Experimental design to evaluate *C. difficile* colonization and disease after colonization with isogenic strains of *H. hepaticus* with differing ability to induce intestinal inflammation in IL-10-deficient mice. IL-10-deficient mice with and without IBD were challenged with spores from *C. difficile* strain VPI 10463 and monitored for up to 31 days for *C. difficile* colonization and clinical severity of disease. B) Lipocalin-2 levels in feces from IL-10−/− mice measured by ELISA at day 14 post-colonization with *H. hepaticus* CDT+ or *H. hepaticus* CDT- or mock-challenged with vehicle (n = 2-6 mice/group). C) Shannon diversity index of cecal and colon contents from mice 14 day after colonization with wild-type *H. hepaticus* (*HhCDT*+) or CDT-deficient H. hepaticus (*HhCDT-*). D) Feces collected over time or E) cecal contents at day 31 post-*C. difficile* spore challenge were plated anaerobically on selective agar plates to quantify *C. difficile* burden. F-G) Mice colonized with *H. hepaticus* CDT+ or *H. hepaticus* CDT-were scored for clinical disease severity and compared to mice mock-challenged with vehicle. ANOVA and Tukey test, ***p<0.001.

Given that the absence of intestinal inflammation is associated with higher gut microbial diversity (Figure 1 and 5C) and resistance to *C. difficile* colonization (Figure 2), we hypothesized that the lack of inflammation in mice colonized with CDT-deficient *H. hepaticus* would result in the preservation of colonization resistance to *C. difficile*. Indeed, cultivatable *C. difficile* was not detected in the feces of SPF IL-10-deficient mice colonized with CDT-deficient *H. hepaticus* at any time point post-*C. difficile* spore challenge (Figure 5D). We confirmed this finding in cecal contents, and found no detectable *C. difficile* colonization in mice at 31 days post-spore challenge (Figure 5E).

Clinical disease severity was evaluated in mice with IBD (*HhCDT+* colonized) or without IBD (*HhCDT−* colonized) following *C. difficile* spore challenge. We observed more severe clinical disease in mice with CDI and underlying IBD compared to mice with IBD that was unaccompanied by comorbid disease (Figure 5F). Mice resistant to colonization by *C. difficile*, including IL-10-deficient mice harboring SPF microbiotas or those additionally colonized with CDT-deficient *H. hepaticus* did not develop IBD (Figure 5D and Figure 5E), and this was reflected in the absence of clinical disease in non-inflamed mice (Figure 5G). As expected, vehicle-challenged mice or mice lacking intestinal inflammation did not develop clinically noticeable signs of disease (Supplemental Figure 2). In summary, these results support the notion that intestinal inflammation is required for *C. difficile* colonization, and that disease severity associated with CDI differs depending on the mode of colonization susceptibility and the presence of underlying comorbid IBD.

## DISCUSSION

While the negative impact of CDI on patients with IBD has been reported in epidemiologic studies (1–3), the specific mediators of IBD-induced susceptibility to *C. difficile* colonization and infection remain elusive. This may be partially due to the lack of a suitable murine model to explicitly dissect the relationship between IBD and CDI, in the absence of antibiotic perturbation of the microbiota. Previous studies using mouse models to explore concurrent IBD and CDI required antibiotic pretreatment to sufficiently render mice susceptible to *C. difficile* colonization and subsequent CDI-associated disease (37–39). However, antibiotic-induced perturbations of the gut microbiota may mask mechanisms specific to IBD-induced *C. difficile* susceptibility and infection. The development of a mammalian model system that does not require antibiotics to disrupt colonization resistance is necessary for the mechanistic investigation of specific IBD-induced mediators of *C. difficile* colonization and infection. Here, we establish a mouse model of comorbid IBD and CDI that demonstrates intestinal inflammation associated with IBD is sufficient to shape a microbiota such that is susceptible to persistent *C. difficile* colonization, obviating the need for antibiotics.

The previously reported studies of CDI in mice induce IBD using dextran sodium sulfate (DSS), a chemical colitogen that damages the intestinal epithelium that results in colitis, require the use of antibiotics for *C. difficile* colonization and subsequent CDI-associated disease (37, 38, 40). While these murine studies have provided key insights in CDI in the context of IBD, including a role for aberrant Th17 responses induced in IBD for increased risk for CDI disease severity (39), the requirement for antibiotic pretreatment to render animals susceptible to *C. difficile* colonization is a caveat that limits use of these models for investigating IBD-specific mediators of *C. difficile* colonization and infection. In our mouse model, IBD results due to a combination of genetic predisposition in the host via IL-10-deficiency and a microbiota trigger through *H. hepaticus* colonization (15). This particular model of IBD is appealing since disease develops through the loss of tolerance to the resident gut microbiota, mimicking human IBD etiology (41, 42). Given that colonization resistance to *C. difficile* is largely mediated by the intestinal microbiota, our mouse model of comorbid IBD and CDI is well suited for investigating the relationship between CDI and the perturbations of the gut microbial community induced by IBD.

Our results demonstrate key differences in *C. difficile* colonization dynamics and CDI-associated disease in mice with IBD compared to non-IBD controls. The antibiotic cefoperazone was used to disrupt the microbiota and demonstrate that *C. difficile* burden in IL-10-deficient mice are similar to that in studies performed in wild-type mice (22). Experiments with IL-10-deficient mice with *H. hepaticus*-triggered intestinal inflammation led to the disruption of the microbiota and revealed that mice with IBD are indeed susceptible to stable *C. difficile* colonization, while genetically identical mice lacking IBD are resistant to colonization. Interestingly, there was a higher frequency of mice colonized with *C. difficile* in the context of cefoperazone pretreatment compared to intestinal inflammation accompanying underlying IBD. The lower rate of *C. difficile* colonization in IBD we observed suggests that intestinal inflammation may stochastically shape the gut microbiota may be stochastic, whereas alterations in gut microbiota following antibiotic treatment is more predictable, given the specificity of these drugs. We have previously shown that in the setting of advanced age, representing a distinct immune- and microbiota-altered host state that increases risk for *C. difficile* infection, CDI results in age-related alterations in neutrophil and eosinophil responses in mice (25). While we found no significant difference in neutrophil responses in mice with IBD-induced or antibiotic-induced CDI, IBD was associated with differences in eosinophil response compared to CDI following antibiotic use. Future work will explore the emerging role of eosinophil responses in intestinal bacterial infection and IBD.

Concurrent CDI in patients with IBD increases morbidity and mortality (7, 43), and this clinical observation has been recapitulated in previous studies in mice demonstrating significantly worse CDI-associated disease following DSS-induced colitis (37–39). While these findings in mice indicated that IBD and antibiotics predisposed animals to develop severe CDI, the specific contribution underlying IBD is unclear in these mouse models due to the requirement for antibiotic pretreatment following colitis-trigger to induce susceptibility to *C. difficile* colonization and infection. Our study explored disease severity in mice with concurrent IBD and antibiotic-independent CDI and also found that these animals developed worse disease compared to mice with IBD alone, albeit not to the degree if disease severity in mice with antibiotic-induced CDI in the absence of underlying IBD. In our model of IBD and CDI, we surprisingly found mice colonized with the virulent toxigenic strain of *C. difficile* yet active toxin was low or undetectable in these mice. This difference in *C. difficile* toxin, rather than bacterial burden, may explain the lack of severe CDI in mice with IBD. Furthermore, IBD-associated CDI was accompanied with delayed weight loss that was not as substantial as antibiotic-associated CDI. Furthermore, comorbid IBD and CDI resulted in less severe clinical disease compared to antibiotic-associated CDI. *C. difficile* strain 10463 used in our studies does not express binary toxin, however it does express toxin A and B (44). Toxin-mediated intestinal damage and pathogenic inflammatory responses mediate CDI-associated disease caused by this highly virulent strain of *C. difficile*. One important limitation of the assay used in this study to quantify *C. difficile* toxin is that this assay does not detect the presence of inactive toxin. Therefore, we cannot rule out the possibility that *C. difficile* toxin is indeed being produced in mice with IBD colonized with *C. difficile*, and that this toxin is not active, and additional studies are needed to follow up this intriguing finding. Regardless, our data suggest key differences in toxin regulation by the intestinal milieu associated with IBD-induced CDI versus antibiotic-induced CDI that warrant further exploration. Taken together, these results indicate key difference between antibiotic-induced versus IBD-induced CDI development and pathogenesis and describe a mouse model of comorbid IBD and CDI.

The significant impact of *C. difficile* infection in the health outcomes of patients with IBD warrants the exploration of mouse models that allows for the systematic study of IBD and concomitant CDI. This study provides a foundation to probe antibiotic-independent mechanisms of *C. difficile* pathogenesis that are specific to the IBD-associated intestinal milieu. It is crucial to elucidate mechanisms of *C. difficile* pathogenesis in the setting of underlying IBD in order to specifically address the clinical problem of CDI in the IBD population. It may not be appropriate to extrapolate mechanisms of susceptibility to *C. difficile* colonization and CDI pathogenesis revealed in antibiotic-induced models to the IBD-induced intestinal milieu and pathogenic synergism of comorbid IBD and CDI. Future studies will aim to distinguish antibiotic-independent structural features of the intestinal microbial community that permit *C. difficile* colonization specifically in the setting of underlying IBD. Ultimately, the use of this model of IBD and CDI will assist in the development and testing of new clinical approaches to predict, diagnose, and manage *C. difficile* infection in the IBD population.

## FIGURE LEGENDS

**Supplemental Figure 1. Intestinal inflammation alters relative abundance of gut microbiota.** Relative abundance of bacterial families in cecal and colon contents from IL-10-deficient mice with IBD (SPF+*Hh*) and without IBD (SPF) at 14 days post-*H. hepaticus* colonization or mock challenge with vehicle.

**Supplemental Figure 2. IL-10-deficient mice not colonized by *C. difficile* lack clinical signs of disease.** Clinical disease severity was assessed following *C. difficile* spore challenge in IL-10−/− mice due to were colonized with CDT-deficient *H. hepaticus* or vehicle 14 days prior to spore challenge, representing various groups of mice without IBD.

## ACKNOWLEDGMENTS

This study was funded by grant U01AI12455 awarded to V.B.Y. by the National Institute of Allergy and Infectious Diseases at the National Institutes of Health. In addition, L.A.C. was supported by grant number UL1TR002240 from the National Center for Advancing Translational Sciences (NCATS).

